# Bin Chicken: targeted metagenomic coassembly for the efficient recovery of novel genomes

**DOI:** 10.1101/2024.11.24.625082

**Authors:** Samuel T. N. Aroney, Rhys J. P. Newell, Gene W. Tyson, Ben J. Woodcroft

## Abstract

Recovery of microbial genomes from metagenomic datasets has provided genomic representation for hundreds of thousands of species from diverse biomes. However, low abundance microorganisms are often missed due to insufficient genomic coverage. Here we present Bin Chicken, an algorithm which substantially improves genome recovery through automated, targeted selection of metagenomes for coassembly based on shared marker gene sequences derived from raw reads. Marker gene sequences that are divergent from known reference genomes can be further prioritised, providing an efficient means of recovering highly novel genomes. Applying Bin Chicken to public metagenomes and coassembling 800 sample-groups recovered 77,562 microbial genomes, including the first genomic representatives of 6 phyla, 41 classes, and 24,028 species. These genomes expand the genomic tree of life and uncover a wealth of novel microbial lineages for further research.

## Introduction

Metagenomics has become an invaluable tool in microbial ecology, and the recovery of thousands of metagenome-assembled genomes (MAGs) from environmental and clinical projects is now common. However, even with these advances, microorganisms lacking genomic representation remain prevalent, particularly in environmental samples^1^. One limitation of standard genome recovery methods is that they often fail to recover low abundance microorganisms due to insufficient sequencing depth^2^. Despite their rarity, low abundance lineages can be critical to microbial community function by performing unique metabolism or by forming a reservoir of genetic diversity critical for restoration after community stress^3,4^. Rare microorganisms often have disproportionate activity, sometimes acting as keystone species (e.g. autotrophic nitrifiers^5^ or methanotrophs^6^) that perform important roles for the ecosystem^7,8^. Due to their lack of genomic representation, much remains to be discovered about low abundance microorganisms.

Deeper sequencing can provide sufficient depth for the recovery of low abundance microorganisms, though at additional cost. An alternative approach is to pool metagenomic reads from multiple samples, coassembling all reads together as if they were from a single sample. Coassembly has the advantage that no further sequencing cost or physical sample manipulation is required. However, pooling reads from multiple samples increases the complexity of assembly, which may result in lower quality genomes^9–13^. The other more practical challenge of coassembly is that assembling combinations of samples can be computationally demanding. Individual coassemblies might require too much RAM, or there may be too many different combinations of samples to coassemble. For instance, even from a small set of 10 metagenomes, there are 120 possible subsets of three samples which can be coassembled.

Metagenome coassembly requires a method to prioritise sample sets for analysis. Sample groupings have previously been made via kmer signatures^14–18^, or through the use of relevant metadata such as host identity, location or other environmental parameters^13,18,19^. Alternatively, entire sample sets can be coassembled at terabase-scale with MetaHipMer^18,20^. These techniques do not prioritise coassemblies based on the novelty or taxonomy of the individual genomes likely to be recovered, making them suboptimal for the recovery of novel genomes.

The use of multiple metagenomes can also improve genome recovery by providing a co-abundance signal useful for binning^21^. These ‘co-binning’ metagenomes can increase genome quality without contributing reads to the assembly^22^. However, even for single-sample assembly, choosing the ideal set of samples to use for co-binning is an unsolved problem. While coassembly and co-binning offer many benefits, the difficulty in choosing the most successful and computationally efficient approach has hampered their general use.

Here, we introduce Bin Chicken, a tool designed to enhance genome recovery in metagenome assembled genomes (MAGs) through three key functions. First, Bin Chicken predicts which coassemblies will yield the greatest novel diversity by identifying highly conserved regions, or “windows,” within marker genes that are shared across metagenome read sets. Second, it leverages this information to recommend sample sets that are ideal for co-binning. Lastly, using taxonomically identified marker genes, Bin Chicken can group samples for coassembly based on specific taxonomic interest—such as taxa underrepresented in reference genome databases—aligning with researchers’ targeted study needs.

## Results

### Bin Chicken guides coassembly by targeting novel genomes

Bin Chicken is a MAG recovery pipeline that maximises recovery of novel diversity by automatically identifying sets of samples for coassembly and co-binning (**Figure 1**). In particular, it identifies groups of samples that share novel marker genes predicted to have sufficient combined coverage for assembly and recovery of their associated genomes (10X minimum, **Figure S2, Supplementary Note 1**). In order to focus on unrecovered diversity, it discards window sequences with less than genus-level divergence from those in reference genomes (**Supplementary Note 1**). Bin Chicken then searches for groups of samples which share the most window sequences, under the hypothesis that these sample groups share the most recoverable diversity. Identical matching of sequence windows is used to reduce the risk of forming chimeric bins from near-relatives (**Supplementary Note 1**).

**Figure 1:**
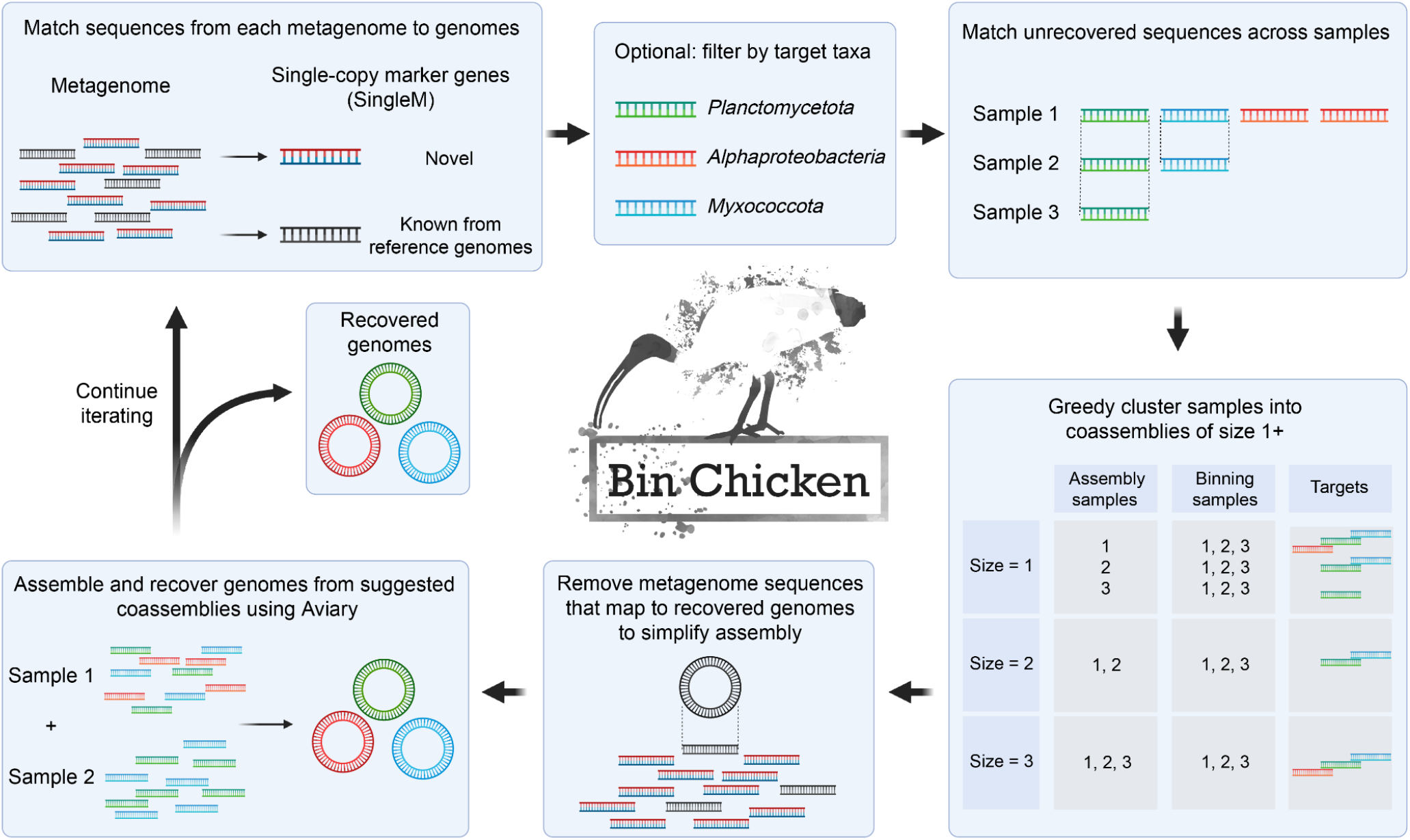
Bin Chicken uses marker genes to choose the best assemblies for metagenome genome recovery. Bin Chicken searches metagenomes and genomes to find single-copy marker genes to match them between metagenomes and genomes and between metagenomes. These links can be filtered by taxonomy to target a particular lineage. The matches are then used to greedily cluster samples into coassemblies of the desired size (1+). Samples for differential coverage binning are also provided by searching the matching sequences for a particular coassembly against all other metagenomes. Metagenomes are also mapped against matching genomes so that the non-mapping sequences can be provided for coassembly. Coassembly and genome recovery is then performed by Aviary^43^. After genome recovery, the entire workflow can be iterated upon to target the remaining novel diversity.

With coassemblies selected, genome recovery begins. To simplify the assembly process, reads from chosen coassemblies are first mapped against reference genomes to allow removal of sequences from already recovered genomes. Samples are coassembled using standard metagenomic assemblers and binning programs (see methods). Samples are chosen for co-binning based on the number of sequences that match each coassembly’s shared window sequences. Once genomes have been recovered, they can inform sample choice for the next iteration of Bin Chicken, ensuring that sequences used for sample grouping remain novel.

By default, Bin Chicken prioritises coassemblies predicted to produce the most recoverable diversity with at least genus-level novelty (**Supplementary Note 1**). However, since marker gene sequences have been assigned taxonomy by SingleM, they can additionally be filtered to focus genome recovery on target taxonomic clades.

### Global coassembly efficiently recovers novel genomes

Bin Chicken was applied to ∼200,000 metagenomes that were publicly available in December 2021^1^. From the >10,000 suggested coassemblies, we chose to assemble those estimated to contain the most recoverable diversity, 433 in total. In addition, we chose 367 coassemblies that targeted 77 of the 93 phyla that were most underrepresented in reference genome databases (<10 genomes in GTDB R214). Coassembling 1,286 samples in 800 groups, Bin Chicken recovered the Rare Biosphere Genomes (hereafter “RBGs”), comprising 77,562 MAGs of at least medium quality^23^ from 38,495 species (**Figure 2, Supplementary Data 2**).

**Figure 2:**
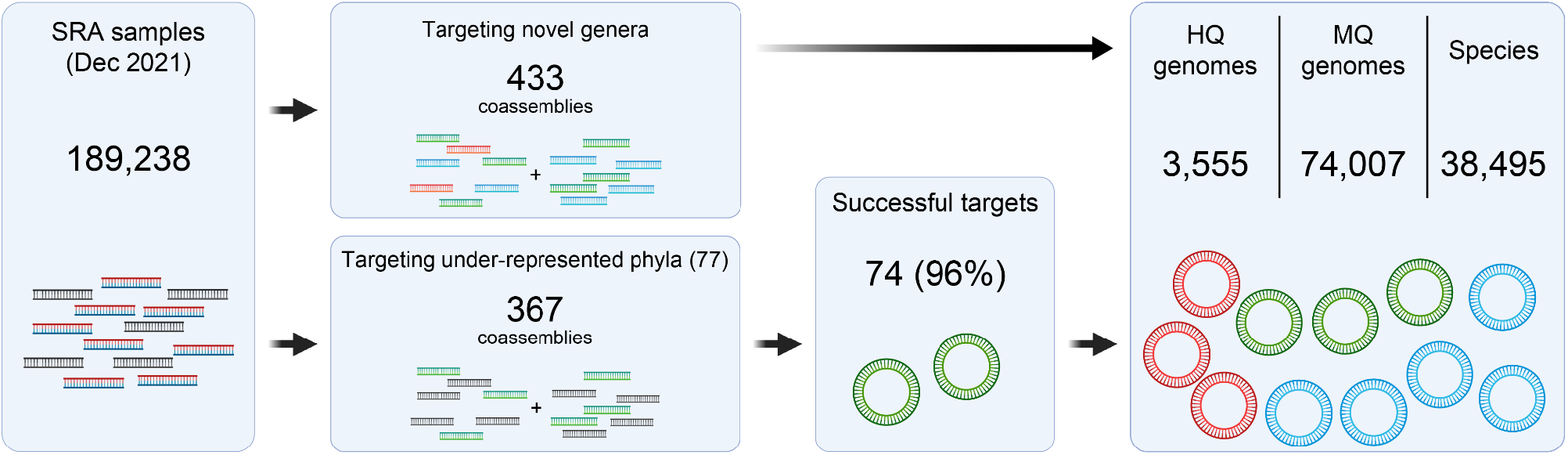
Recovering novel genomes from public metagenomes. Coassemblies were chosen by iteratively running Bin Chicken, targeting recoverable novel diversity and under-represented phyla with <10 genomic representatives. A target was considered successful if Bin Chicken recovered a matching genome from the coassembly targeting a given phyla. SRA - sequence read archive; HQ - high-quality genome according to MIMAG standards; MQ - medium quality genome.

Of the 93 underrepresented phyla, 77 had sufficient coverage for Bin Chicken to suggest coassembly. These coassemblies recovered genomes of novel species in 73 of these phyla (96%), confirming Bin Chicken’s ability to recover new genomes from underrepresented lineages (**Supplementary Note 4**). In 8 of these phyla, genomes from novel classes were recovered.

To address concerns about the quality and chimerism of RBGs, we randomly selected 30 coassemblies, and compared them to the same samples assembled individually. The co-binning step was kept consistent to specifically study differences introduced by co-assembly. Coassembly produced 49% more species-level genome bins on average, with few genomes recovered exclusively by single-sample assembly (9.9%). Where the same species was recovered by both methods, the quality of the coassembled genome was 2.8% higher (completeness – 5 × contamination, p<0.001, n=2,170, ANOVA), and produced no discernible increase in genome chimerism (**Figure S3, Supplementary Note 2**). Amongst the total set of RBGs, the rate of chimerism (9.8%) was also comparable to the rate observed in reference MAGs in the Genome Taxonomy DataBase (GTDB; 8.0%)^12,24^. These findings suggest that coassembly does not inherently reduce genome quality or increase chimerism as might be hypothesized, but instead results in higher quality genome bins than single-sample assembly. However, as previously observed^13^, it was very rare that multiple genomes from the same species were recovered from the same coassembly (15 out of the 77k genomes), suggesting that coassembly may interfere with the recovery of strain-specific genotypes.

### Rare Biosphere Genome novelty

The taxonomic novelty of RBGs was established against a baseline of genomes from GTDB (version R220) and several recent large-scale genome recovery efforts^25–30^. *De novo* tree inference revealed that addition of the RBGs increased the known phylogenetic diversity of Bacteria by 12% and Archaea by 18% (with 12% and 22% species growth, respectively, **Figure 3A,B**). The known phylogenetic diversity was increased by >25% in 35 phyla (**Supplementary Table 5, Supplementary Table 6**). Among the RBGs were the first bacterial representatives of 6 phyla, 38 classes, 215 orders, 1,299 families, 8,993 genera and 21,787 species (**Figure 3A, Supplementary Note 5**), and the first archaeal representatives of 3 classes, 26 orders, 175 families, 1,051 genera and 2,241 species (**Figure 3B**). Together, the RBGs represent a substantial advance in our genomic understanding of the microbial tree of life.

**Figure 3:**
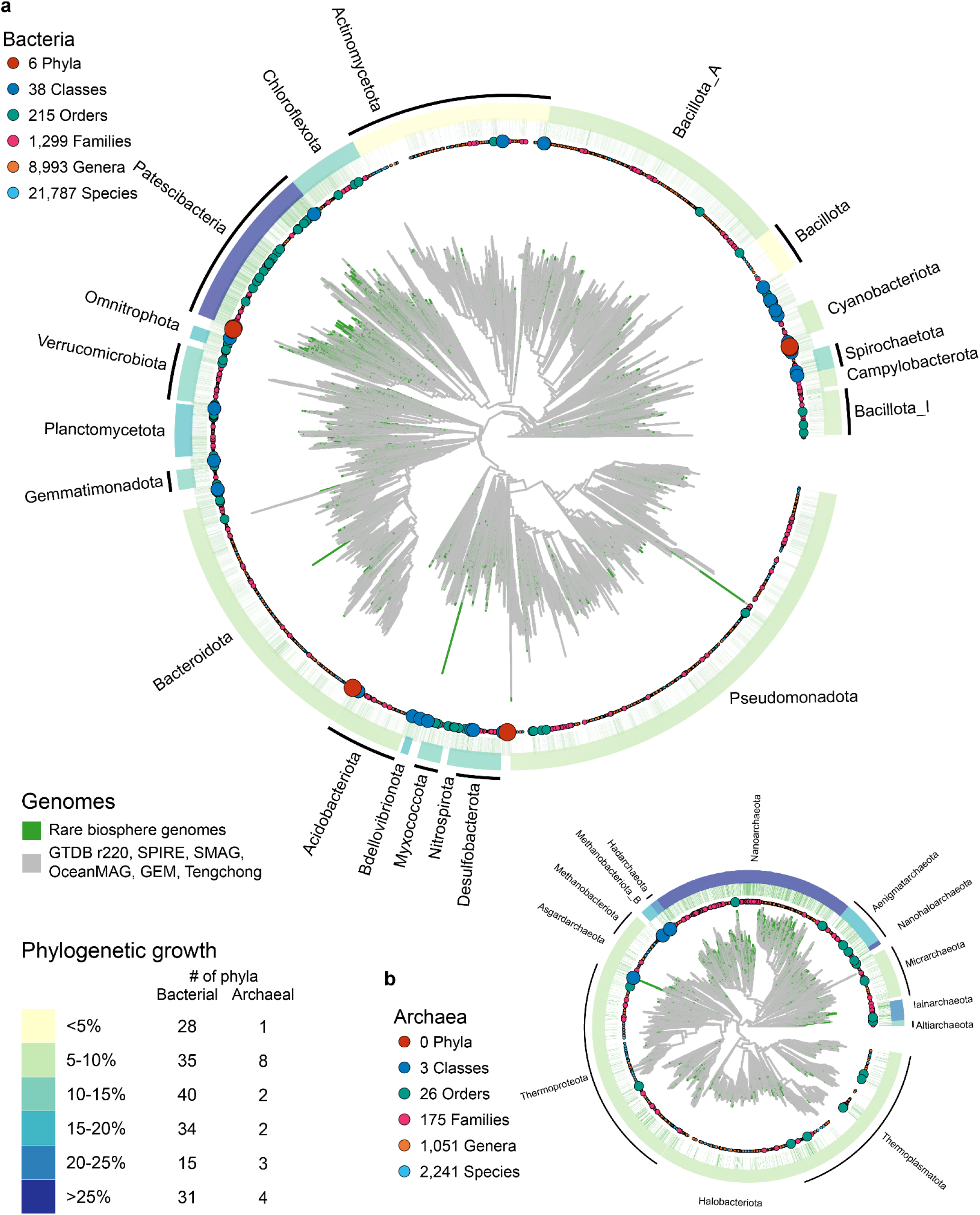
Coassembly recovers genomic novelty across the prokaryotic tree of life. Genomic phylogenetic trees of (a) bacterial and (b) archaeal genomes from Rare Biosphere Genomes (RBGs), GTDB R220, SPIRE, SMAG, OceanMAG, GEM, Tengchong hot springs and UHGG. Branches are coloured by exclusivity to RBG. Circles represent RBG with species-to phyla-level novelty. Inner green band is coloured for RBG. Outer band is coloured by the phylogenetic growth of RBG for the most diverse phyla. GTDB - the Genome Taxonomy Database; SPIRE - Searchable, Planetary-scale mIcrobiome REsource; SMAG - soil MAGs; GEM - Genomes from Earth’s Microbiomes; UHGG - the Unified Human Gastrointestinal Genome.

To establish the efficiency of Bin Chicken in recovering novel species, we compared it with SPIRE, the largest scale effort at MAG recovery from public metagenomes to date^30^. Relative to efforts previous and GTDB R214, SPIRE recovered 83k novel species from single-sample assembly and single-sample binning of 99,146 metagenomes (∼500 Tbp). RBG, in contrast, contains 24k novel species relative to all previous efforts, recovered from only 800 coassemblies of 1,286 samples (17.8 Tbp) relative to an updated reference genome set which includes SPIRE (**Supplementary Table 2, Supplementary Table 3, Supplementary Table 4, Supplementary Data 3**). Thus, Bin Chicken recovered >40X the number of novel species per assembled metagenome (1.3% of samples, 29% of the novel species in SPIRE). We conclude that Bin Chicken provides an efficient means of recovering novel species diversity.

On average, 30 novel species were recovered per coassembly (three archaea and 27 bacteria). Accumulation curves of novel species were linear, strongly suggesting that much untapped genomic diversity recoverable by Bin Chicken is present in publicly available metagenomes (**Figure 4A**). Accumulation curves at higher taxonomic ranks have begun to taper (**Figure 4B, Figure S7**), particularly at phyla level, but they have not yet plateaued suggesting many high-level lineages remain unrecovered (**Figure 4C**).

**Figure 4:**
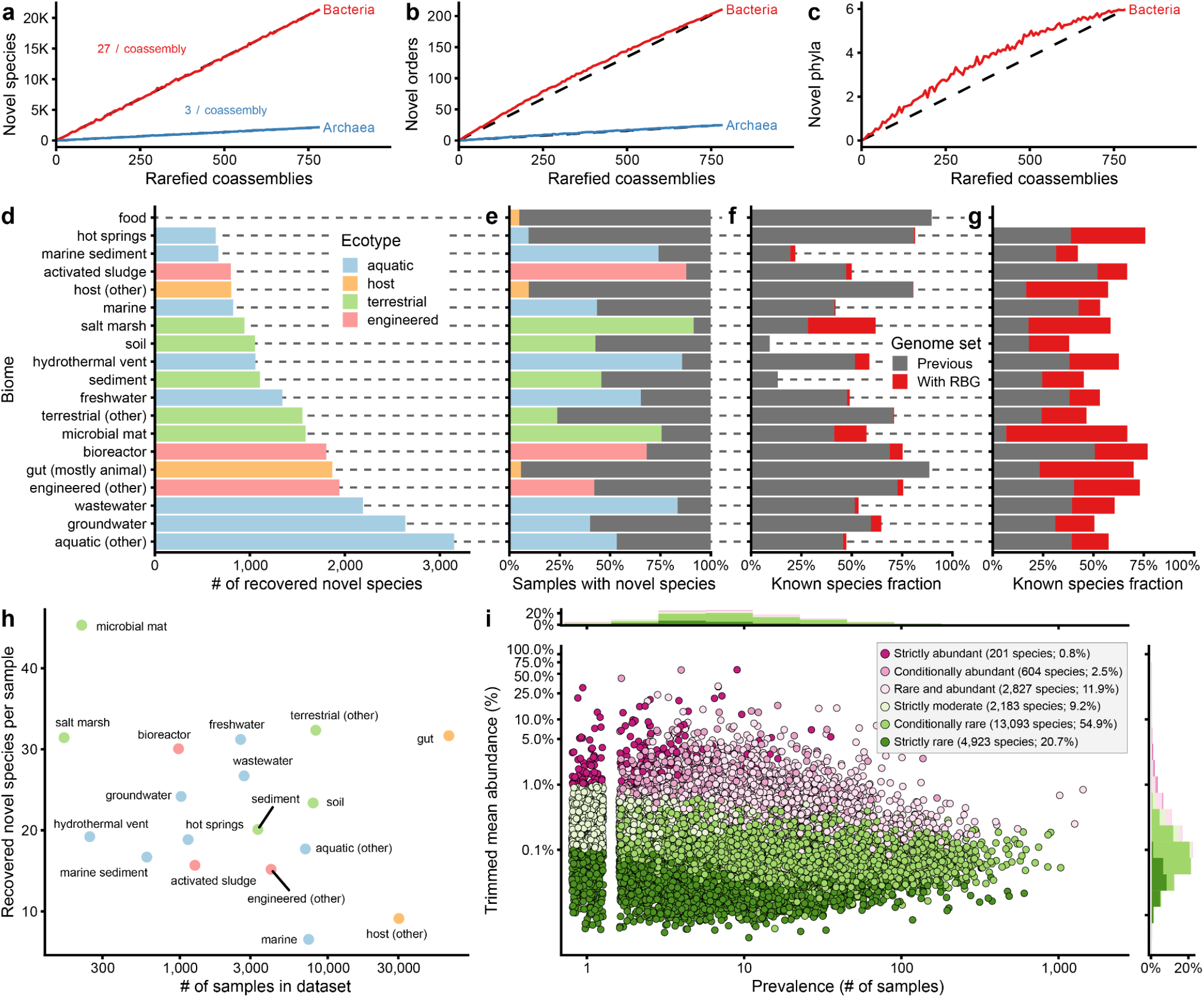
Rare Biosphere Genomes were found from biomes worldwide. Accumulation curves of rarefied coassemblies against counts of novel Rare Biosphere Genome (RBG) species (a), orders (b) and phyla (c), averaged across 100 iterations. Dashed line added for linear comparison. (d) Source biome (host or environment) for novel RBG species. Bar colour represents sample ecotype. (e) Proportion of samples containing matching novel RBG species single-copy marker genes. (f, g) Median improvement in known species fraction based on single-copy marker genes, abundance weighted, for (f) all samples and (g) source samples. (h) Recovered novel RBG species per sample analysed against the number of samples for each biome in the dataset. (i) Abundance vs prevalence of novel RBG species across samples. Estimated through single-copy marker genes. Point colour represents the abundance category, determined using a 0.1% cutoff for rarity and a 1.0% cutoff for abundance. Bars represent binned density of genomes for x (above) and y (right) axes. Category selection required >95% of samples to meet the respective cutoff. 197 novel species were not assigned to a rarity group due to being undetected.

### Abundance and prevalence of rare taxa

Novel RBG species were recovered from nearly all major biomes, particularly aquatic (3,151 species, 12% of recovered species), groundwater (2,636, 10%) and wastewater (2,191, 8%) (**Figure 4D, Figure S6A, Supplementary Data 5, Supplementary Data 6**). The prevalence and abundance of novel RBG species was estimated by adding them to the SingleM reference database and re-estimating community profiles of the public metagenome dataset. Novel RBG were present in 35k/180k samples, notably in most aquatic and engineered samples (53% and 55%), many terrestrial samples (36%) but were missing from most host samples (7%; **Figure 4E, Figure S6B**). In addition, while novel species were found across 424 different biomes, 43% were only found within 1 biome and 89% within at most 5 different biomes. Novel RBGs increased the median fraction of each microbial community that was represented at the species level (the “known species fraction”) across biomes by an average of 4% and by 27% among samples that were used for coassembly (**Figure 4F,G, Figure S6C**,**D**). For instance, the median known species fraction of microbial mat metagenomes increased by 16% across all 217 public samples and by 60% across the 35 samples used for coassembly. When normalised by the number of samples used for coassembly, the most prolific biomes were microbial mats, salt marshes, freshwater and other terrestrial environments (**Figure 4H**).

We categorized the novel RBG species by how rare they were based on their global distribution of abundances, with a cutoff of <0.1% for “rare” and >1% for “abundant” as per Dai et al. ^31^. Species were classified as either strictly rare (<0.1% in >95% of samples), strictly abundant (>1% in >95% of samples), conditionally rare/abundant (sometimes rare/abundant but never the opposite), strictly moderate (never rare/abundant) or rare and abundant (remaining species). Within public samples, novel RBG tended to be rare, with more than 75% classified as conditionally or strictly rare, never present at more than 1% abundance (**Figure 4I**). Indeed, 90% (21,676) of novel RBG species were present at <1% abundance in the samples from which they were recovered. This was expected since the standard single-sample assembly genome recovery is biased towards abundant lineages, thus the remaining un-recovered species are likely rarer, with insufficient coverage for single-sample recovery. The rare biosphere remains largely unexplored, but the RBG represents a significant step towards understanding and characterising the rare microbiome and opens the door to further recovery of novel genomes through large-scale coassembly.

### Metabolic capacity of the rare microbiome

We clustered 596 million proteins identified from all known genomes into 210 million clusters with minimum sequence identity of 40%. Similar to novel species recovery, accumulation curves of protein homology groups found only in novel RBG species were near-linear, with 21 million new protein clusters discovered at a rate of 868 per species and 26,105 per co-assembly (**Figure 5A,C**). Bin Chicken recovered 33% of the amount of novel protein clusters in SPIRE from only 1.3% of the samples. Bin Chicken thus provides an efficient means of recovering novel proteins in addition to novel species. Much of the microbial protein universe remains undiscovered or not associated with a genome^32^.

**Figure 5:**
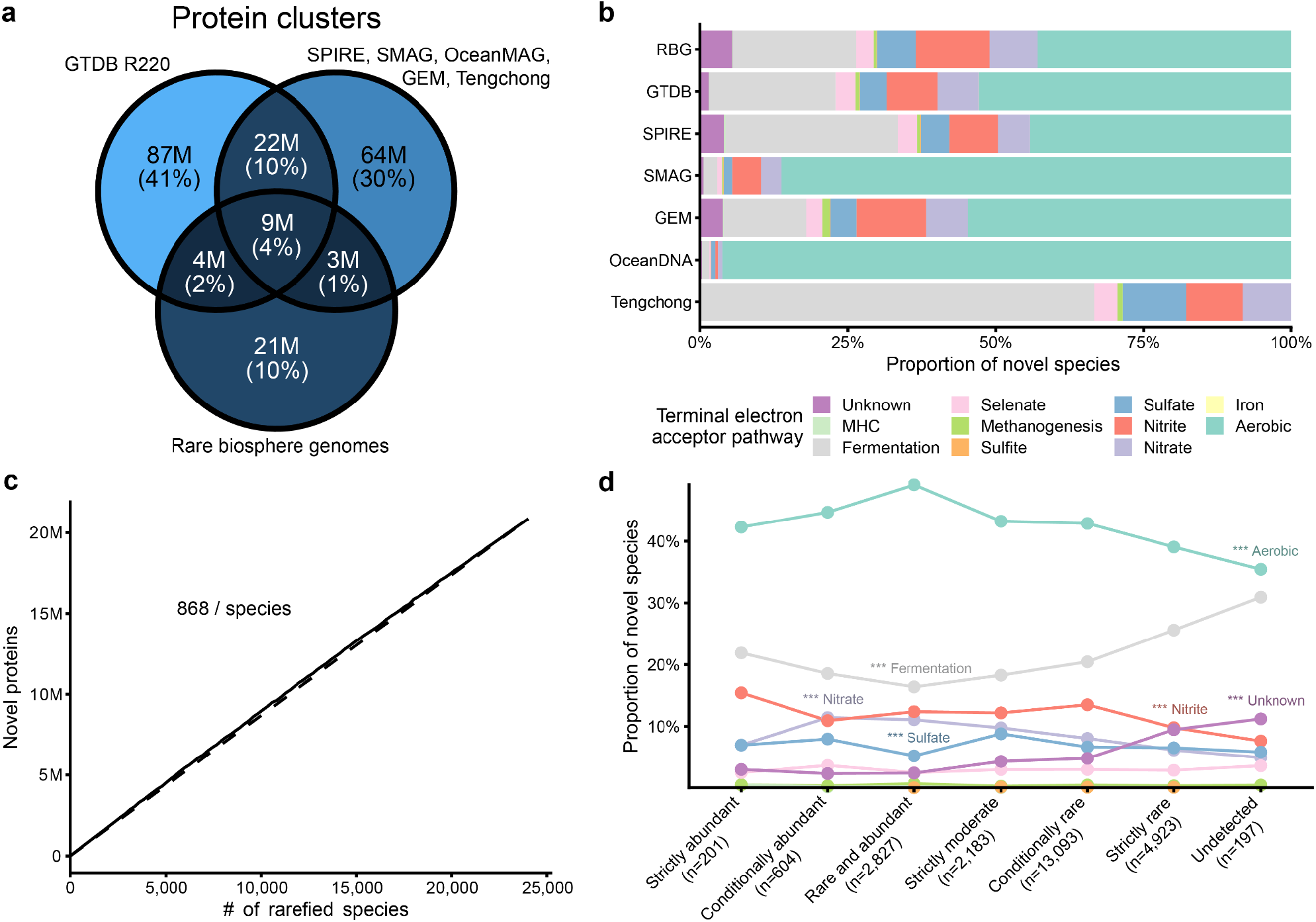
Metabolic capacity and protein clusters of Rare Biosphere Genomes. (a) Venn diagram of protein clusters between species representatives from GTDB R220, SPIRE, SMAG, OceanMAG, GEM, Tengchong, UHGG and the Rare Biosphere Genomes (RBGs). (b) Proportion of novel species that could be assigned to major terminal electron acceptor pathways. Genomes with multiple assignments were assigned to the most thermodynamically favourable pathway. (c) Accumulation curve of rarefied RBG species against counts of novel RBG protein clusters, averaged across 5 iterations. Dotted line added for linear comparison. (d) Proportion of novel RBG species split into their abundance rarity categories with pathway assignment from above. n: number of species within that category. ANOVA with binomial distribution and false-discovery rate correction found significantly changing pathways, *** p<0.001. GTDB - the Genome Taxonomy Database; SPIRE - Searchable, Planetary-scale mIcrobiome REsource; SMAG - soil MAGs; GEM - Genomes from Earth’s Microbiomes; UHGG - the Unified Human Gastrointestinal Genome.

Annotation of genes with putative functions allowed each novel genome to be categorised according to its most energetically favourable terminal electron acceptor. MAGs recovered by the OceanDNA effort had the largest proportion of aerobic genomes (96%) while RBG had the smallest (43%) except for the targeted MAG recovery from the largely anaerobic Tengchong hot spring sediment samples (0% of 267 novel species) (**Figure 5B**). RBG also had the highest proportion of novel species with unknown metabolism (5.4%), compared to 1.5% in GTDB.

The metabolism of novel RBG species varied significantly with their abundance (**Figure 5D**). Rare species tended to be less aerobic (ANOVA, n=24,028, p=2×10^−15^), possibly due to aerobic organisms having a selective advantage in energy generation, resulting in their greater abundance ^33^. Conversely, fermenters and metabolically unclassified species were more common among rare species (ANOVA, n=24,028, p=1×10^−24^ and p=3×10^−48^). This enrichment of unknown metabolisms suggests that rarer species may perform functions substantially different to those of more abundant species, and to those which grow readily under culture conditions.

### Metabolism of novel phyla

The RBG contains 10 genomes that are the first representatives of six bacterial phyla, almost all of which appear to be anaerobic (**Figure 6, Supplementary Table 7**). Three novel aquatic phyla (**Figure 6A**) were recovered from marine, groundwater and hydrothermal vent samples^34–39^. These genomes encoded various anaerobic metabolisms, including sulfate reduction, nitrate reduction and fermentation. Multi-haem *c*-type cytochromes (MHCs), which may act on a number of potential metabolites, were also observed. The terrestrial novel phylum (**Figure 6B**) was recovered from a salt marsh in the United States. The single genome representing this phylum was predicted to be anaerobic and encoded homoacetogenesis and fermentation pathways. Two novel phyla were recovered from engineered systems (activated sludge, wastewater and bioreactor, **Figure 6C**)^40,41^. The two genomes recovered from the first of these phyla (binchicken_b4) were anaerobic with sulfate reduction and fermentation pathways. In contrast, the two genomes from the other phylum (binchicken_b5) were not predicted strongly as either aerobic or anaerobic, possibly indicating that they are facultative anaerobes. These genomes encoded nitrate reduction, fermentation, and MHCs, but only encoded the TCA cycle, glyoxylate cycle and electron transport chain component V among potential aerobic pathways. All phyla were notably rare in both prevalence and abundance, with 9/10 genomes classified as strictly or conditionally rare and all phyla appearing in only 3-110 samples out of ∼200,000.

**Figure 6:**
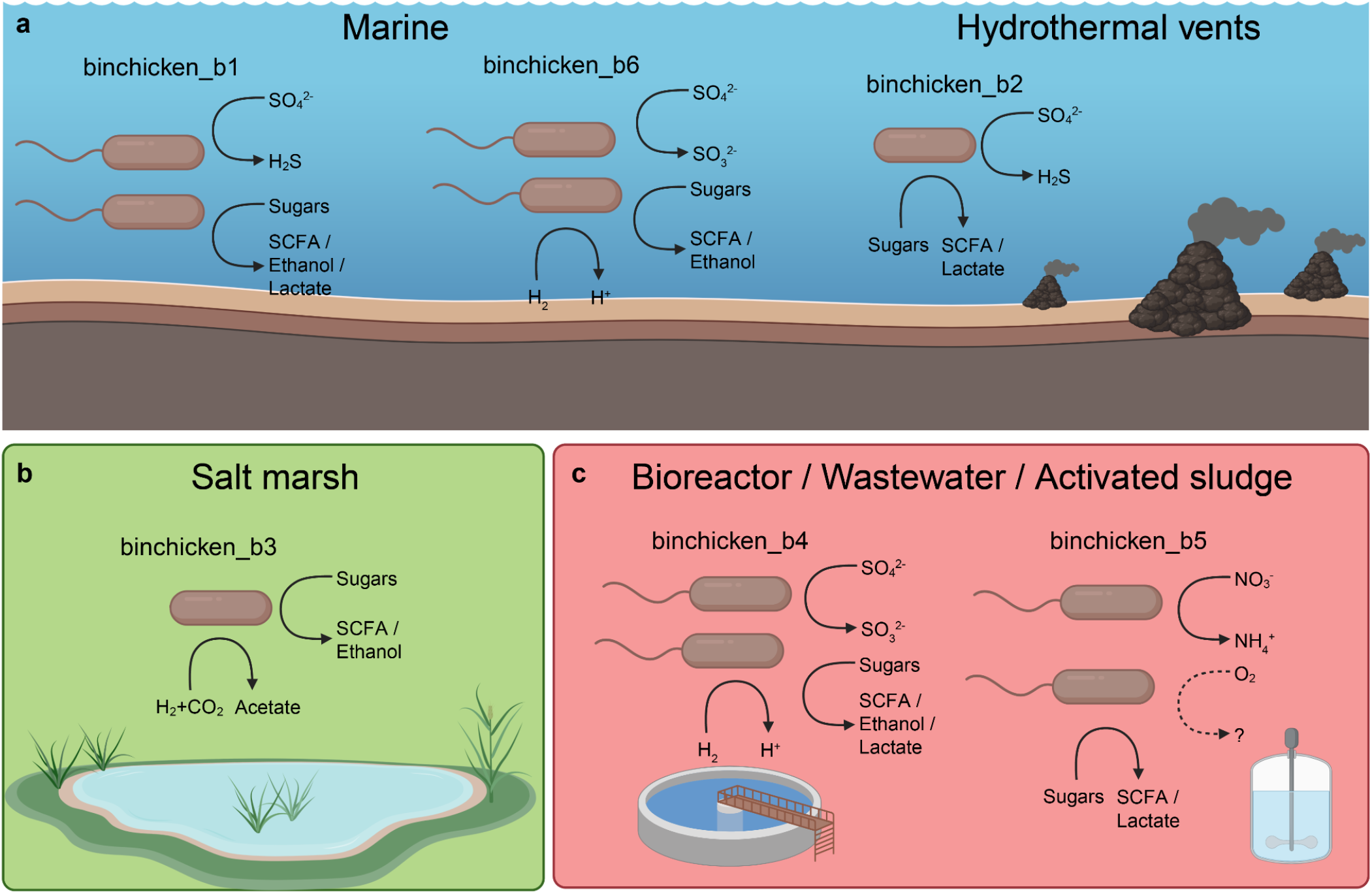
Encoded metabolism of novel phyla. Each cartoon microbe represents an independent genome recovered from the novel phyla indicated above it. Genomes were recovered from aquatic (a), terrestrial (b) and engineered (c) ecotypes. SCFA: short-chain fatty acids.

## Discussion

Metagenomic analysis typically relies on the processing of individual samples, though genome recovery is often limited by insufficient sequencing depth^20^. Here we show that standard metagenomic assembly methods applied to rationally chosen sets of samples markedly improves the quality and quantity of recovered MAGs. Using Bin Chicken, we recovered MAGs from 24k novel species using publicly available metagenomic datasets. These species were prevalent, found in one fifth of all samples, including half of aquatic and engineered samples and a third of terrestrial samples (**Figure 3, Figure 4**). They were also mostly rare, present at <1% relative abundance in most samples where they were detected (76% of species).

The RBGs grow our genomic knowledge of the microbial tree of life appreciably, including the first representation from seven new phylum-level lineages. RBGs were recovered at a much faster rate per sample than other recent large-scale MAG recovery efforts (**Supplementary Table 4**), and the rate of novel lineage discovery remains high even at the phylum level (**Figure 4A-C, Figure S7**). This suggests that continued application of Bin Chicken will efficiently recover further novel diversity.

Despite substantially improved MAG recovery rates from coassembled samples (median 19% improvement, **Figure 4G**), the quantity of known species within public metagenomes remains low, particularly among environmental samples (terrestrial median 18%, aquatic 48%, engineered 68% and host 87%, **Figure S6**). If estimates^42^ that Earth hosts 10^11^-10^12^ microbial species are correct, then >99.99% of species lack genomic representation since <10^7^ are present in databases. Huge quantities of bacterial and archaeal genomic diversity are yet to be discovered, particularly in the rare biosphere, providing an important aim for future work.

## Acknowledgements

S.T.N.A. was supported by the EMERGE National Science Foundation (NSF) Biology Integration Institute (#2022070) and Genomic Science Program of the United States Department of Energy (DOE) Office of Biological and Environmental Research (BER), grants DE-SC0004632, DE-SC0010580 and DE-SC0016440. B.J.W. was supported by Australian Research Council grants (#DP230101171, #FT210100521). We thank those who have provided comments including Andy Leu, Pierre-Alain Chaumeil and Phil Hugenholtz. We thank high performance computing system administrators Chris Williams, Mitchel Haring and Hamish McDonald.

## Author Contributions

S.T.N.A. developed the Bin Chicken program with assistance from R.J.P.N., under the supervision of B.J.W. and G.W.T.. S.T.N.A. applied Bin Chicken to public metagenomes and analysed the recovered genomes. S.T.N.A. and B.J.W. wrote the manuscript with input from G.W.T. and R.J.P.N.. All authors reviewed and approved the final version of the manuscript.

