## Supplementary material for "Bin Chicken: targeted metagenomic coassembly for the efficient recovery of novel genomes": Online methods

### Bin Chicken software

Bin Chicken (github.com/AroneyS/binchicken) is a python wrapper around a Snakemake[1] pipeline that proposes coassembly sample-groups based on the co-occurrence of single-copy marker genes, excluding those genes present in reference genomes (e.g. previously recovered genomes). The Polars Python library[2] was used for data manipulation. Samples were greedily clustered based on exact matching of marker gene sequences with each cluster containing samples with the highest number of matches among samples not yet clustered. Samples for differential abundance binning were chosen based on the number of matches to the co-occurring sequences found in all cluster samples. Sample sequences were then optionally mapped to the reference genomes that were matched to each sample, with only non-mapping sequences being assembled. Assembly and genome recovery were performed using the Aviary pipeline[3].

### Global coassembly survey

Coassemblies were chosen based on the sample-groups proposed by Bin Chicken v0.10.0 using Sandpiper SingleM marker gene lists from all ≥1Gbp SRA metagenomes with at least 1 marker-gene hit, as of 11/2022. SingleM v0.16.0[4] was used with the GTDB R214 metapackage v3.2.0[5]. Bin Chicken was run independently for each SRA study using an 86% divergence cutoff (roughly equivalent to genus-level novelty) against GTDB R214, SPIRE[6], SMAG[7], OceanDNA[8], UHGG v2[9] and GEM[10] genomes and searching for 2-5 sample coassemblies with 20 recovery samples each (more details in **Supplementary Note 3**). Samples were downloaded using Kingfisher v0.3.0[11]. Aviary assembly and genome recovery was run either on the non-mapping (<99% identity or <99% alignment) sequences of coassemblies with the most novel diversity or on the entire sequences of coassemblies targeting phyla with at most 10 representative genomes in GTDB R214. 20 samples were used for co-binning for each coassembly, including the samples that were assembled. Each iteration's recovered genomes were used as reference genomes for the next iteration.

### Metagenomic assembly and genome recovery

Metagenomic assembly was performed using the "aviary assemble" subcommand of Aviary v0.8.2[3]. This includes trimming and quality filtering of reads using fastp v0.23.4[12], followed by assembly by either Metaspades v3.15.4[13] or, if it failed due to insufficient memory (maximum of 500 GB provided), MEGAHIT v1.2.9[14].

Metagenomic genome recovery was performed using the "aviary recover" subcommand from the Aviary pipeline. This includes initial read mapping through CoverM v0.6.1[15] using minimap2 v2.18[16] and subsequent binning by MetaBAT v2.15[17], MetaBAT2 v2.15[18], VAMB v3.0.2[19] and SemiBin2 v2.0.2[20]. Pilot coassemblies also used CONCOCT v1.1.0[21], MaxBin2 v2.2.7[22] and Rosella v0.4.2[23] binners. Bins from all binners were collected and deduplicated using DASTool v1.1.2[24].

### Genome properties and functional annotation

Genomes analysis used a Snakemake workflow (see post_processing/post_processing.smk in github.com/AroneyS/binchicken-genome-processing). Chloroplast genomes were removed when identified. CheckM2 v1.0.2[25] was used to estimate genome completeness and redundant contamination. CoverM v0.6.1[15] was used to estimate the proportion of cross-sample chimeric contigs, by calculating read coverage breadth for each individual sample used for a given coassembly. GUNC v1.0.5[26] was used to estimate genome non-redundant contamination. Genomes with quality (completeness – 5 × contamination) at least 50 were clustered at species-level (95% ANI) using Galah v0.4.0[27]. GTDBtk v2.3.0[28] was used to taxonomically classify MQ genomes.

Gene transcripts and protein sequences were predicted using prodigal v2.6.3[29]. 5S, 16S and 23S rRNA genes were predicted by barrnap v0.9[30]. tRNA genes were predicted by tRNAscan-SE v.2.0.12[31]. All genomes were functionally annotated using eggNOG-mapper v2.1.11 or v2.1.14 using eggNOG DB v5.0.2[32,33]. Database searches for gene sequences were performed using Diamond v2.1.9[34] and MMseqs2 v15.6f452[35]. In addition, all genomes from novel phyla were functionally annotated using DRAM v0.15.0[36]. HydDB[37] was used to further classify identified hydrogenases. Fermentation pathways were manually curated based on KEGG pathways[38], requiring all key reactions. Putative multi-haem *c*-type cytochromes were identified using both Pfam domains for cytochrome *c* and multiple CXXCH-motifs.

Proteins predicted above (609,622,874) were clustered using MMseqs2 v15.6f452[35] using `mmseqs easy-linclust` with `--min-seq-id 0.4 --threads 64` to form 196,305,965 clusters. Rarefaction curves were calculated by sampling 100 steps across the 600M proteins for 5 iterations and averaging the number of unique protein clusters at each step. Genome rarefaction curves were calculated the same way, but with 10-100 iterations and averaging the number of unique novel clades at each step across the coassemblies.

### Coassembly benchmarking

Thirty randomly selected coassemblies (15 from each coassembly strategy) were compared to the samples assembled individually. Genomes were clustered together at species-level (95% ANI) as above. Coassembly increase in species-level bins was calculated as the average across coassemblies of the number of species only found by coassembly divided by the total number of species found by single sample assembly. The number of genomes recovered exclusively by single sample assembly was calculated as the average across coassemblies of the number of species only found by single sample assembly divided by the total number of species found by single sample assembly. Genome quality was compared for species that had genomes recovered by both single sample assembly and coassembly. Where single sample assembly recovered multiple genomes from the same species, the one with the highest quality was used for comparison.

### Phylogenetic and taxonomic diversity analysis

*de novo* genome trees for bacterial and archaeal genomes from novel species (determined based on 95% ANI clustering) were generated using GTDBtk v2.3.0 de_novo_wf[28]. Phylogenetic diversity and gain (additional branch length from Rare Biosphere Genomes) were calculated using GenomeTreeTk v0.1.8[39]. Novel clades were identified through a custom algorithm based on relative evolutionary divergence (RED), similar to that described in Supplementary Figure S1 from Chaumeil et al. [40]. Firstly, RED values were calculated by PhyloRank v0.1.12 (github.com/dparks1134/phyloRank) using the *de novo* genome trees from GTDBtk. Next, the RED values and genome trees were processed using a custom script (see post_processing/name_clades.py in github.com/AroneyS/binchicken-genome-processing). Genomes were sequentially assigned to either pre-existing or novel clades, ordered first by publication order and then by genome quality. For each genome, this involved traveling up the tree until a node was reached within the RED-bounds of a new taxa level (e.g. phylum, class, order, etc.) for which the genome has not yet been assigned. Parents and children of the node were checked to find if any were closer to the median RED of that taxa level (from GTDB R220) than the current node. If the current node was the closest node, it is annotated as a new clade, with the current genome being assigned as the genome representative. This continued until a node was reached that had an offspring from GTDB (or until reaching a previously labeled node). The remaining taxonomy was assigned based on the median taxonomy of the GTDB offspring from that node.

IQ-TREE v2.0.3[41] was used to create more robust genome trees using a representative from each order. Amino acid alignments from GTDBtk de_novo_wf were used as inputs. Ultrafast bootstrap approximation was applied with 1000 replicates (-B 1000). Genomes with phyla and class-level novelty that branched within another phyla or class were reclassified within that phyla or class.

### Prevalence and abundance analysis

Genomes from SPIRE[6], SMAG[7], OceanDNA[8], UHGG v2[9], Tengchong hot springs[42] and GEM[10], with and without Rare Biosphere Genomes were added to the GTDB R220 SingleM metapackage v4.3.0[43] using `singlem supplement`. Taxonomy assignments of OTU tables from all SRA samples were updated with both new metapackages using `singlem renew`. Known species fraction was calculated as the proportion of abundances assigned to species level in community profiles generated from `singlem condense`. In addition, novel RBG species were classified into abundance-based categories as done previously[44], using 0.1% cutoff for rare taxa and 1% cutoff for abundant taxa, except that category selection required >95% of samples to meet the respective cutoff. So, strictly abundant required an abundance of ≥1% in at least 95% of samples with the species present and strictly rare required an abundance of ≤0.1% in at least 95% of samples with the species present. Conditionally abundant/rare had some samples meeting the abundant/rare cutoff but did not meet the criteria for strictly abundant/rare, but also without having more than 5% of samples meeting the opposite cutoff. Strictly moderate had at least 95% of samples at neither ≥1% nor ≤0.1%. Finally, rare and abundant was assigned to remaining species with some presence in the samples.

### Data analysis

R Statistical Software v4.3.1[45] was used for data analysis and figure generation using tidyverse v2.0.0[46], cowplot v1.1.1[47], ape v5.7-1[48] and ggtree v3.9.0.001[49] packages. GNU Parallel v20231122 was used for internal processing[50]. Plotting scripts were run in reproducible Conda environments (github.com/AroneyS/binchicken-genome-processing).

### Data availability

All genomes, metabolic annotations and metadata were deposited on Sharepoint (https://connectqutedu.sharepoint.com/:f:/s/BinChickensupplementarydata/ErJbGzlvzglMoLSAIjv3dtIBaEmqdZXoUHYRQlYYbtSY5Q?e=aIvFdH). Representative genomes from novel species were submitted to ENA in project PRJEB82487.
