## Supplementary notes, figures, tables for "Bin Chicken: targeted metagenomic coassembly for the efficient recovery of novel genomes"

### Supplementary notes, figures and tables for "Bin Chicken: targeted metagenomic coassembly for the efficient recovery of novel genomes"

#### Supplementary Note 1

##### Matching marker gene sequences between samples and between samples and reference genomes

Marker gene sequences between samples and reference genomes were matched to ignore sequences from already recovered genomes. To choose sequence identity cutoffs for different taxonomic ranks, we compared the average nucleotide identity between sequences with known taxonomic divergence within the GTDB R207 SingleM metapackage v3.1.0[1]. We decided to prioritize genomes at primarily genus-level novelty (or above) for this project, so chose a 86% identity cutoff (**Figure S1**). Note that since window sequences are only 60 bp, cutoffs are only meaningful in 1/60th intervals. Also note that this cutoff is imperfect, excluding some relationships with family-level taxonomic divergence and including some relationships with genus-level taxonomic divergence. Cutoffs at higher taxonomic divergence had even more overlap, and were not considered.

Marker gene sequences between samples were matched based on an identical aligned window sequence. We wanted to restrict matching to as specific as possible to mitigate the risk of forming chimeric bins from near-relatives. Since species-level divergence between these sequences is 2bp on average[2], genomes sharing identical window sequences are usually derived from strains of the same species.

##### Choosing the coverage cutoff for window sequences

Bin Chicken uses a coverage cutoff to determine which marker gene window sequences are likely to represent a recoverable genome. A cutoff of 10X coverage was chosen *a priori* based on prior experience with taxonomically-targeted genome recovery[2]. An *a posteriori* analysis proved this cutoff to be an apt choice, with recovery rate below 10X at 24.1% and above at 83.2% (**Figure S2**). When restricted to novel sequences above 10X, the rate fell to 52%.

#### Supplementary Note 2

##### Benchmarking genomes recovered from coassembly against single-sample

We benchmarked Bin Chicken coassembly against single-sample assembly using the same assembly and binning methods. Thirty randomly selected coassemblies (15 from each coassembly strategy style, see **Supplementary Table 1**) were compared to the 70 samples assembled individually. We recovered an average of 49% more species from coassembly than from single-sample assembly (**Figure S3A**). For species found by both coassembly and single-sample, the genome recovered from coassembly had on average 2.8% higher genome quality, 2.7% higher genome completeness, 0.15% higher contamination, unchanged contig chimerism (contig-covered fraction from a single sample) and 0.9% switched from GUNC-fail to pass (**Figure S3B-F**). This indicates that coassembled genomes were better quality, with a low proportion of contig chimerism and fewer chimeric genomes than from single-sample assembly.

### Supplementary Note 3

##### Iterative coassembly of public metagenomes with Bin Chicken

We applied Bin Chicken to 186,155 public metagenomes from NCBI SRA using SingleM Sandpiper marker gene data[2], in a batched, iterative manner, progressively choosing coassemblies that were likely to yield novel species compared with genomes known at the beginning of the iteration (**Supplementary Table 1, Supplementary Data 1**). Three quarters (75%, 29,007/38,495) of RBG species are novel relative to these reference genomes. Coassemblies were run as a concatenation of the non-mapping (<99% identity or <99% alignment) sequences of chosen samples. We also ran coassemblies targeting phyla with low representation, for which the whole of each sample was concatenated for assembly. Not all attempted coassemblies were completed successfully, mostly due to finding that the SRA samples were single-ended reads (these were skipped, **Supplementary Data 4**).

### Supplementary Note 4

##### Rare Biosphere Genome properties

Iterative coassembly produced 77,562 (65,150 bacterial and 8,857 archaeal) Rare biosphere genome (RBG) MAGs of at least medium quality, including 3,555 (3,000 bacterial and 555 archaeal) high-quality MAGs according to MIMAG standards[3] (**Figure S4, Figure S5**). These genomes formed 38,495 (34,610 bacterial and 3,889 archaeal) 95% ANI genomospecies clusters.

##### Targeting phyla with low genomic representation successfully recovered more representatives

In addition to choosing coassemblies based on diversity, we also targeted phyla with low genomic representation in GTDB R214 (<10 genomes; **Figure 2**). Of the 93 phyla that meet this criterion, 77 had a coassembly with at least 2 matching sequences (a very low cutoff compared to the expected 35-37 marker genes per genome). By running these coassemblies, we recovered genomes from 74 of these phyla, including 8 novel classes (against GTDB R214). This indicates that not only does the targeting produce novel genomes, but it can also target unrecovered genomes with sequences greatly divergent to known sequences (i.e. class-level divergence).

#### Supplementary Note 5

##### Taxonomic assignment of novel species

Novel RBG species were assigned taxonomy based on *de novo* genome trees formed from GTDB R220 representative genomes, in addition to novel species from UHGG, GEM, OceanDNA, SPIRE, SMAG and Tengchong. Genomes were sequentially assigned to either pre-existing or novel clades through a custom algorithm based on relative evolutionary divergence (RED), similar to that described in Supplementary Figure S1 from Chaumeil et al. [4]. See Methods for more details.

More robust genome trees were generated using IQ-TREE, with only a single representative from each class (including novel classes) and, independently, from each order (including novel orders). Each novel phyla, class and order was examined to check for incongruencies between taxonomic assignment and tree structure, i.e. to find if a given clade caused another clade to become paraphyletic. Only 8/215 bacterial and 2/26 archaeal Bin Chicken clades with at least order-level novelty branched with strong bootstrap support (>80%) within a different phyla or class than their taxonomy assignment. These clades, including 1 putative novel phyla and 9 putative novel classes, were manually assigned appropriate taxonomy deferring to the IQ-TREE trees.

### Supplementary Figures

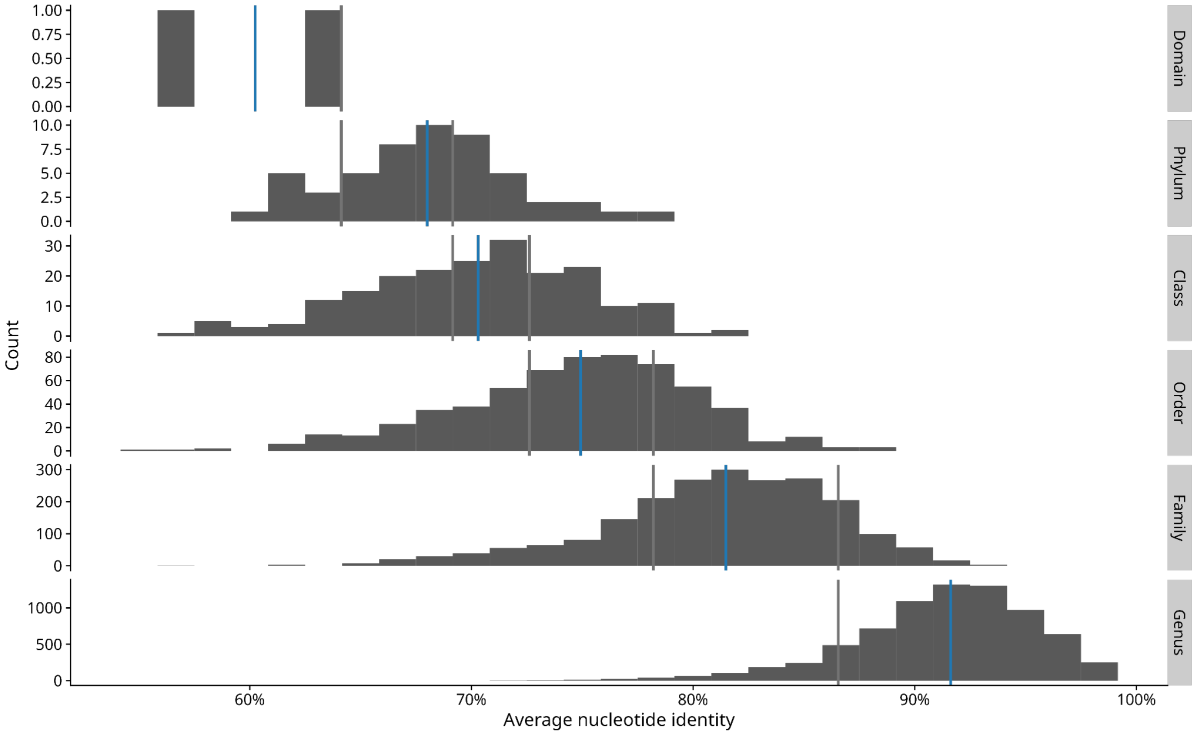


**Supplementary Figure 1: Average nucleotide identity between SingleM window sequences with varying taxonomic divergence.** Each comparison represents the average of the possible comparisons between genomes within the chosen taxonomy. Blue lines represent mean identity for each population and gray lines represent the midpoint between their means.

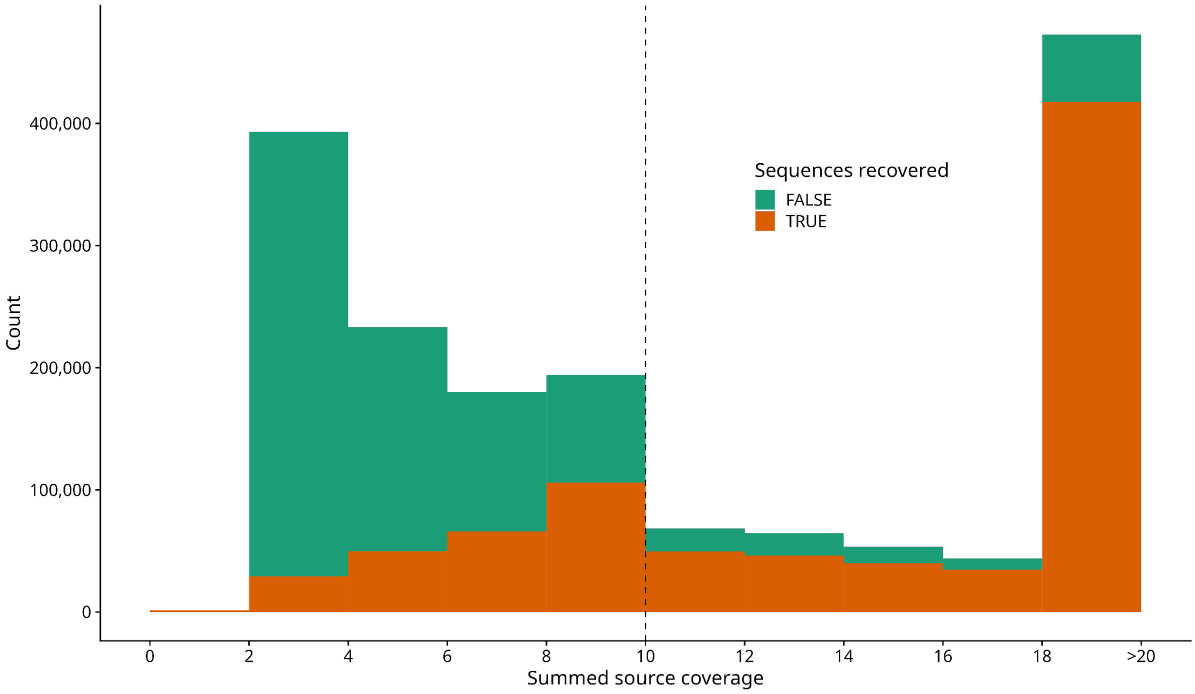


**Supplementary Figure 2: Recovery of marker gene sequences (SingleM) compared to summed source coverage.** The estimated coverage from marker gene sequences found in all source samples per coassembly was summed. Each sequence was matched to sequences found in genomes recovered from the same coassembly.

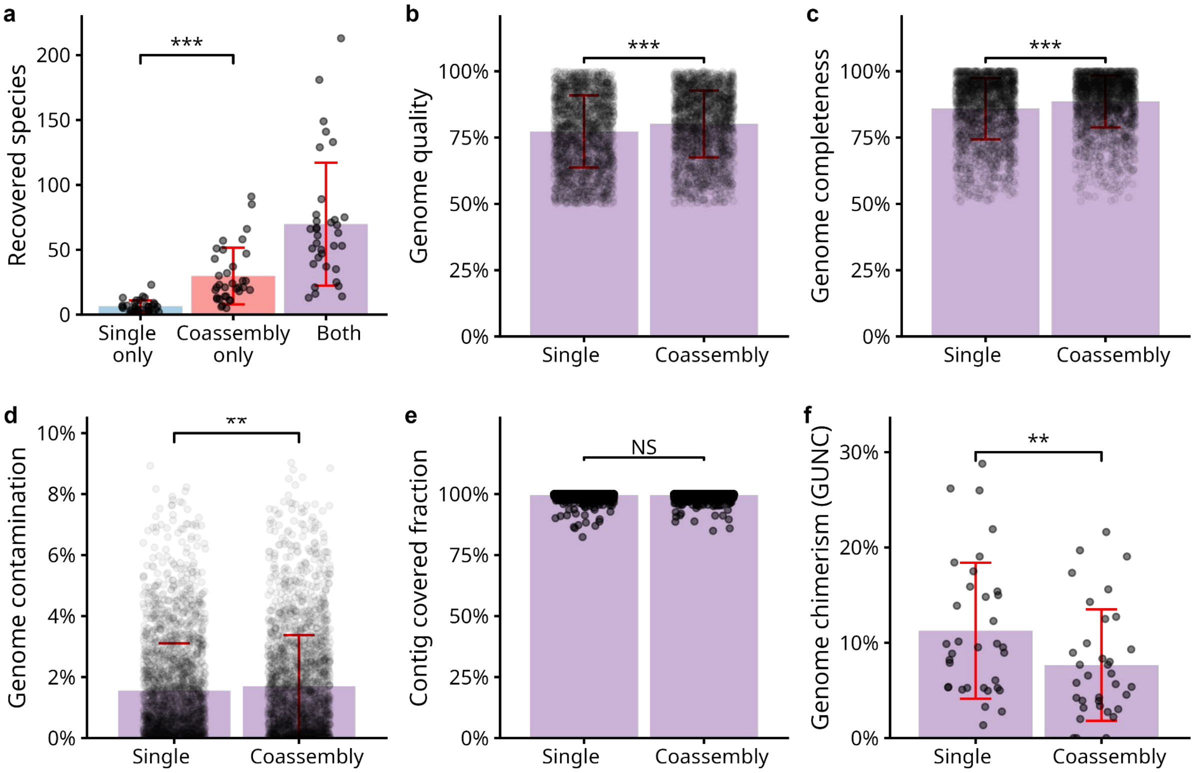


**Supplementary Figure 3: Benchmarking genomes recovered from coassembly against single-sample assembly.** (a) Comparison of genomes recovered from 30 randomly selected coassemblies and 70 single-sample assemblies from each of the samples that were coassembled. (b,c,d,e,f) Comparison of genomes recovered from coassembly and single-sample from within 95% ANI species clusters. Columns represent mean ± standard deviation of (b) genome quality (CheckM2 completeness – 5 × contamination), (c) completeness, (d) contamination, (e) length-weighted contig covered fraction from one sample and (f) proportion of genomes with GUNC detected chimerism per sample-set. Points represent genomes (b, c, d, e) or sample-sets (f). Data were modeled using ANOVA. NS: not significant, ** p<0.01, *** p<0.001.

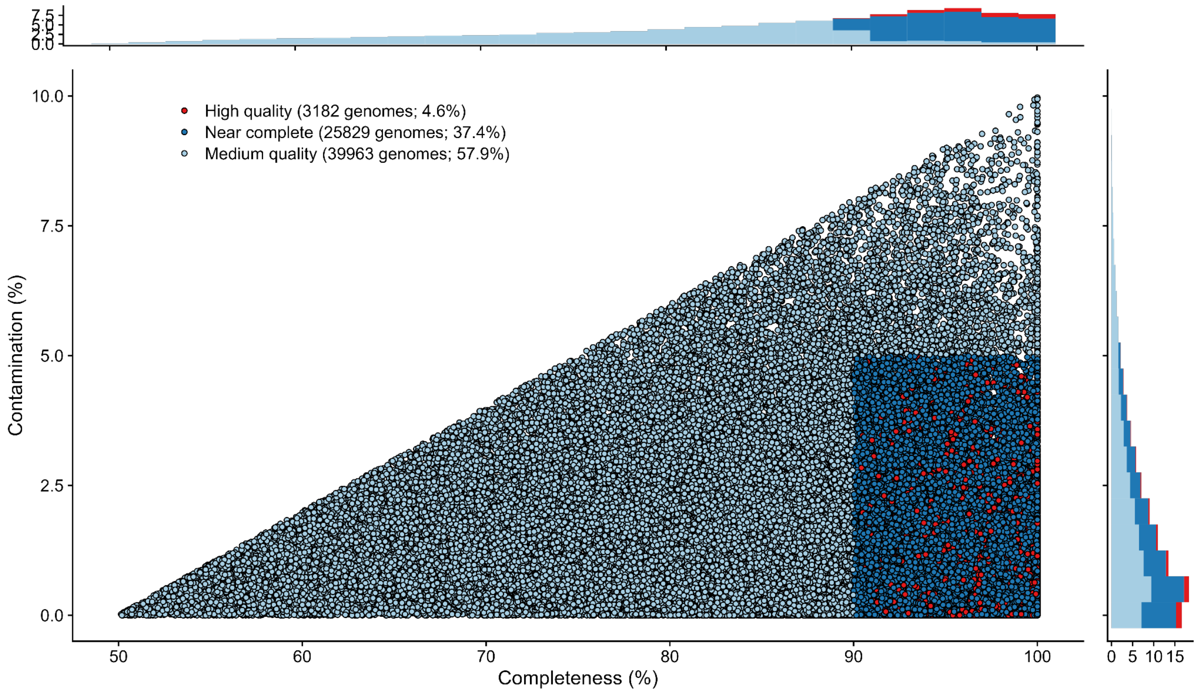


**Supplementary Figure 4: Genome completeness and contamination of Rare Biosphere Genomes.** Completeness and contamination were estimated using CheckM2. Genomes were classified as high-quality if they were >90% complete and <5% contaminated in addition to rRNA and tRNA cutoffs.

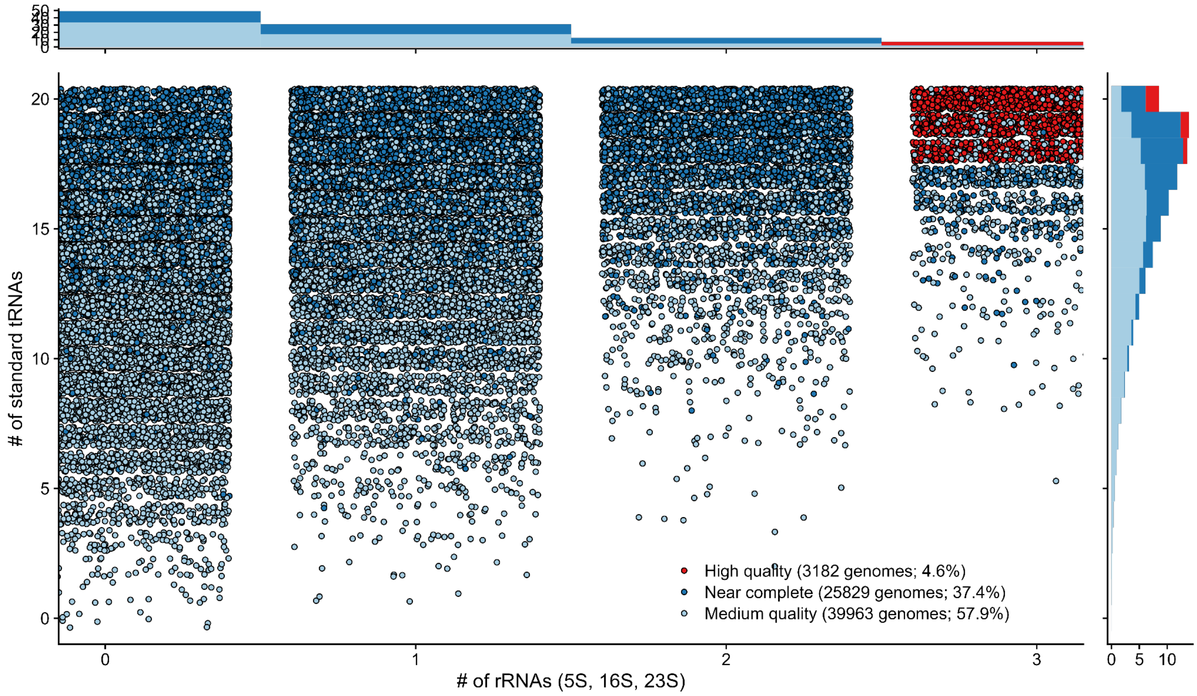


**Supplementary Figure 5: Genome rRNA and tRNA counts for Rare Biosphere Genomes.** Genomes were classified as high-quality if they had each of the three rRNAs and at least 18/20 standard tRNAs in addition to genome quality cutoffs.

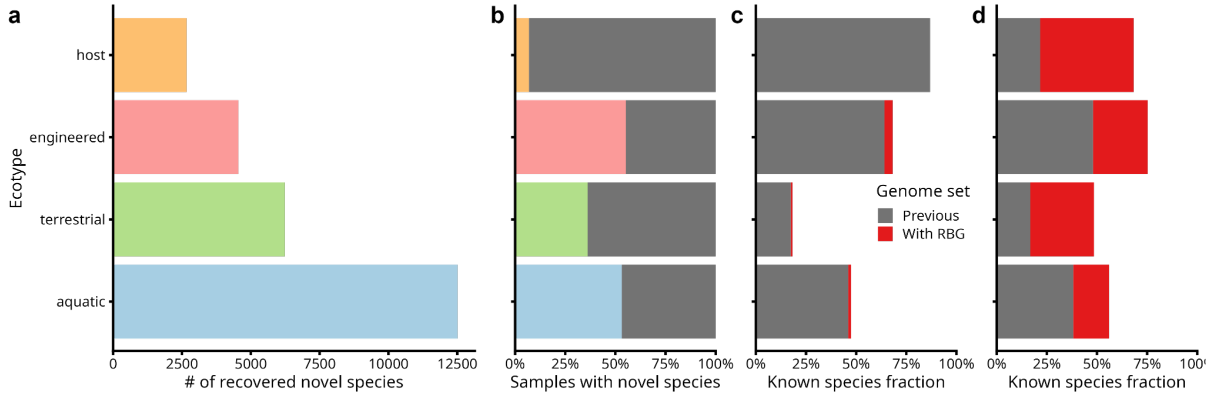


**Supplementary Figure 6: Rare biosphere genomes source and prevalence summarized by ecotype.** (a) Source sample ecotype for novel RBG species. (b) Proportion of samples containing matching novel RBG species single-copy marker genes (SingleM). (c, d) Median improvement in known species fraction based on single-copy marker genes (SingleM), abundance weighted, for (c) all samples and (d) coassembled samples.

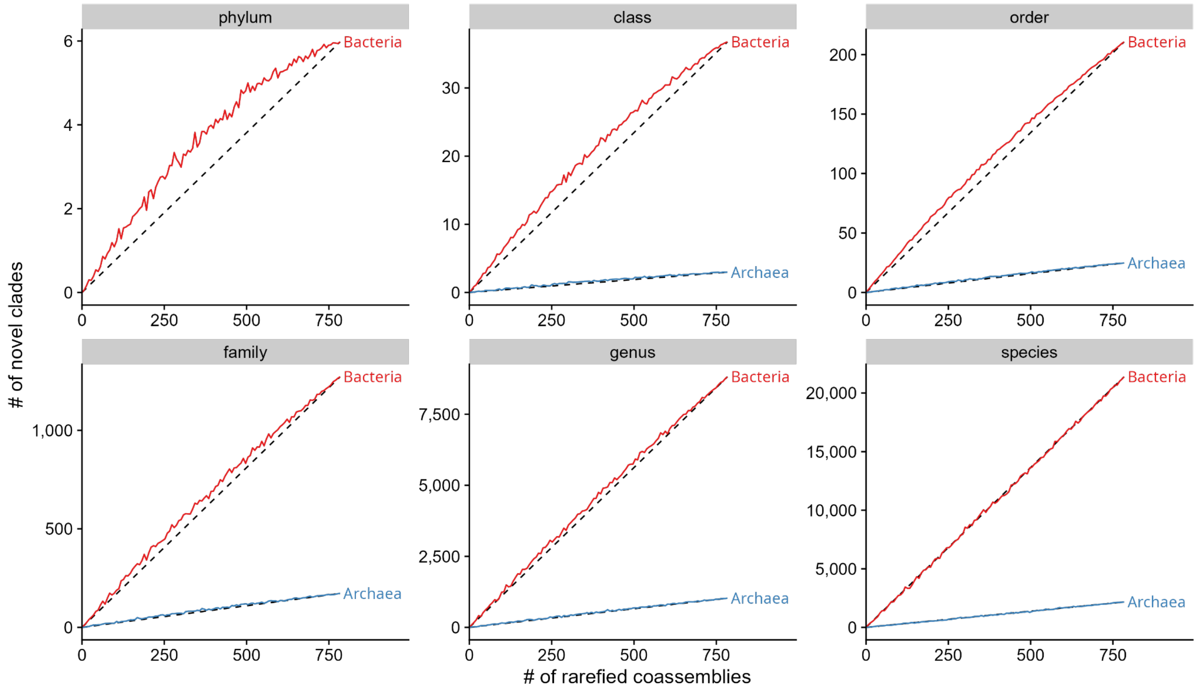


**Supplementary Figure 7: Novel clade coassembly accumulation curves.** Coassemblies were rarefied over 100 iterations, and the number of novel clades was averaged across iterations.

#### Supplementary Table 1

Summary of Bin Chicken coassembly strategy. In general, each run set iterated on the genomes recovered from the previous set to target new genomes. Run set 4 also included genomes from UHGG, GEM, OceanDNA, SPIRE, SMAG and Tengchong. “Target” style refers to coassemblies chosen to target each phyla from GTDB R214 with fewer than 10 genome representatives. The final target run was to target novel phyla from SPIRE, GEM, SMAG, OceanDNA and previously recovered Rare Biosphere Genomes. “Diversity” style refers to coassemblies chosen for having the most recoverable diversity.

| **Style** | **Run set** | **# of coassemblies (completed)** | **# of samples per coassembly** |
|---|---|---|---|
| Target | 1 | 49 (46) | 2 |
| Diversity | 1 | 10 (10) | 2 |
| Target | 2 | 10 (9) | 2-3 |
| Diversity | 2 | 14 (14) | 2 |
| Target | 3 | 71 (67) | 2-3 |
| Diversity | 3 | 100 (99) | 2 |
| Target | 4 | 38 (38) | 2-4 |
| Diversity | 4 | 179 (157) | 2-3 |
| Target | 5 | 112 (110) | 2-5 |
| Diversity | 5 | 170 (153) | 2-4 |
| Target | 6 | 105 (97) | 2-5 |

Supplementary Table 2

Archaeal genome novelty per dataset against GTDB R220 genomes. Final column indicates Bin Chicken novelty against GTDB R220 genomes, ignoring genomes from the other MAG-sets. Novel clades were assigned in publication order (UHGG, GEM, OceanDNA, SPIRE, SMAG, Tengchong and Bin Chicken). Values are inclusive (i.e. a novel phyla is also a novel class, etc.).

| | **UHGG** | **GEM** | **OceanDNA** | **SPIRE** | **SMAG** | **Tengchong** | **Bin Chicken** | **Total** | **Bin Chicken Against R220** |
|---|---|---|---|---|---|---|---|---|---|
| **Phylum** | | | | | | | | | |
| **Class** | | 2 | | 5 | | | 3 | 12 | 10 |
| **Order** | | 7 | 2 | 28 | | 2 | 26 | 65 | 45 |
| **Family** | | 41 | 11 | 217 | 11 | 25 | 175 | 480 | 316 |
| **Genus** | | 160 | 60 | 1,217 | 110 | 125 | 1,051 | 2,723 | 1,598 |
| **Species** | 1 | 371 | 231 | 2,838 | 473 | 267 | 2,241 | 6,421 | 2,926 |

Supplementary Table 3

Bacterial genome novelty per dataset against GTDB R220 genomes. Final column indicates Bin Chicken novelty against GTDB R220 genomes, ignoring genomes from the other MAG-sets. Novel clades were assigned in publication order (UHGG, GEM, OceanDNA, SPIRE, SMAG, Tengchong and Bin Chicken). Values are inclusive (i.e. a novel phyla is also a novel class, etc.).

| | UHGG | GEM | OceanDNA | SPIRE | SMAG | Bin Chicken | Total | Bin Chicken Against R220 |
|---|---|---|---|---|---|---|---|---|
| **Phylum** | | 2 | 1 | 17 | | 6 | 27 | 15 |
| **Class** | | 16 | 2 | 69 | 4 | 38 | 136 | 91 |
| **Order** | | 75 | 6 | 317 | 35 | 215 | 648 | 392 |
| **Family** | 1 | 340 | 52 | 1,927 | 241 | 1,299 | 3,859 | 2,167 |
| **Genus** | 15 | 1,935 | 532 | 15,530 | 3,247 | 8,993 | 30,237 | 13,268 |
| **Species** | 54 | 4,556 | 2,442 | 53,225 | 13,309 | 21,787 | 95,319 | 27,408 |

#### Supplementary Table 4

Microbial novelty per sample used for genome recovery. Novelty for both archaea and bacteria compared against GTDB R220 genomes. Novel clades were assigned in publication order (UHGG, GEM, OceanDNA, SPIRE, SMAG, Tengchong and Bin Chicken).

| MAGset (# of samples) | | **Bin Chicken** (1,286) | **SPIRE** (99,146) | **GEM** (10,450) | **OceanDNA** (2,057) | **SMAG** (3,304) | **Tengchong** (152) |
|---|---|---|---|---|---|---|---|
| **Novel lineages per 1000 samples** | **Phylum** | 5 | <1 | <1 | <1 | | |
| | **Class** | 32 | <1 | 2 | <1 | 1 | |
| | **Order** | 187 | 3 | 8 | 4 | 11 | 13 |
| | **Family** | 1,146 | 22 | 37 | 31 | 76 | 164 |
| | **Genus** | 7,810 | 169 | 200 | 288 | 1,016 | 822 |
| | **Species** | 18,684 | 565 | 471 | 1,299 | 4,171 | 1,757 |

#### Supplementary Table 5

Archaeal phylogenetic growth through addition of Rare Biosphere Genomes to *de novo* species genome tree generated from GTDB R220, UHGG, GEM, OceanDNA, SPIRE, SMAG and Tengchong genomes. Abbreviations: # prior - Number of species in previous genome databases; # RBG - Number of Rare Biosphere Genome species; PG - phylogenetic growth; SG - species growth.

| **Clade** | **# prior** | **# RBG** | **PG%** | **SG%** |
|---|---|---|---|---|
| d__Archaea | 10054 | 2246 | 17.6 | 22.3 |
| p__JACRDV01 | 1 | 1 | 53.6 | 100 |
| p__Nanoarchaeota | 1686 | 973 | 28.6 | 57.7 |
| p__Nanohaloarchaeota | 39 | 20 | 28.1 | 51.3 |
| p__Undinarchaeota | 9 | 6 | 26.3 | 66.7 |
| p__Iainarchaeota | 185 | 87 | 24.5 | 47 |
| p__SpSt-1190 | 15 | 6 | 24 | 40 |
| p__Hadarchaeota | 66 | 25 | 23.8 | 37.9 |
| p__Aenigmatarchaeota | 436 | 149 | 18.7 | 34.2 |
| p__Altiarchaeota | 65 | 15 | 14.3 | 23.1 |
| p__Hydrothermarchaeota | 31 | 7 | 13.4 | 22.6 |
| p__Methanobacteriota_B | 140 | 12 | 13.2 | 8.6 |
| p__Thermoproteota | 2415 | 359 | 10 | 14.9 |
| p__Asgardarchaeota | 356 | 61 | 9.8 | 17.1 |
| p__Thermoplasmatota | 1899 | 217 | 9.7 | 11.4 |
| p__Halobacteriota | 1700 | 190 | 9.1 | 11.2 |
| p__Micrarchaeota | 563 | 82 | 9 | 14.6 |
| p__EX4484-52 | 38 | 2 | 8.6 | 5.3 |
| p__B1Sed10-29 | 38 | 4 | 6.4 | 10.5 |
| p__Methanobacteriota | 319 | 24 | 5.5 | 7.5 |
| p__Methanobacteriota_A | 37 | 0 | 0 | 0 |

#### Supplementary Table 6

Bacterial phylogenetic growth through addition of Rare Biosphere Genomes to *de novo* species genome tree generated from GTDB R220, UHGG, GEM, OceanDNA, SPIRE and SMAG genomes. Includes the top 19 phyla with >50 total genomes. Abbreviations: # prior - Number of species in previous genome databases; # RBG - Number of Rare Biosphere Genome species; PG - phylogenetic growth; SG - species growth.

| **Clade** | **# prior** | **# RBG** | **PG%** | **SG%** |
|---|---|---|---|---|
| d__Bacteria | 180852 | 21808 | 12.3 | 12.1 |
| p__Patescibacteria | 11144 | 5195 | 26.7 | 46.6 |
| p__UBA10199 | 90 | 34 | 22.6 | 37.8 |
| p__QNDG01 | 41 | 18 | 22.5 | 43.9 |
| p__Sumerlaeota | 73 | 28 | 22.2 | 38.4 |
| p__Dependentiae | 268 | 102 | 21.6 | 38.1 |
| p__Riflebacteria | 44 | 15 | 20.3 | 34.1 |
| p__Desulfobacterota_G | 79 | 18 | 18.9 | 22.8 |
| p__Hydrogenedentota | 124 | 37 | 18.5 | 29.8 |
| p__Krumholzibacteriota | 171 | 61 | 18.2 | 35.7 |
| p__Calditrichota | 80 | 23 | 17.8 | 28.7 |
| p__Omnitrophota | 1067 | 306 | 17.6 | 28.7 |
| p__Zixibacteria | 278 | 94 | 17.4 | 33.8 |
| p__Electryoneota | 73 | 25 | 17.3 | 34.2 |
| p__UBP6 | 55 | 8 | 17.2 | 14.5 |
| p__Bdellovibrionota | 645 | 152 | 17 | 23.6 |
| p__Bacillota_F | 77 | 22 | 16.8 | 28.6 |
| p__Eisenbacteria | 185 | 41 | 16.4 | 22.2 |
| p__Margulisbacteria | 167 | 34 | 16.1 | 20.4 |
| p__KSB1 | 117 | 39 | 15.9 | 33.3 |

### Supplementary Table 7

Novel phyla habitat and metabolic capacity

| Phyla (habitats) | Genome | Stats | Habitat/s | Metabolism |
|---|---|---|---|---|
| binchicken_b1<br><br>Marine (16)<br>Seawater (7)<br>Groundwater (3)<br>Biofilm (1) | SRP200020_co2_210 | **Completeness:** 96%<br>**Contamination:** 0.7%<br>**Assembly size:** 4.6Mbp<br>**# contigs:** 289<br>**GC:** 35%<br>**rRNAs:** -<br>**tRNAs:** 18 | ***Source***<br>Marine<br>***Marker-gene***<br>Marine (15)<br>Seawater (7)<br>***Rarity***<br>Strictly rare | ***Aerobicity***<br>Predicted anaerobic (99.9% confidence)<br>***Energy***<br>Glycolysis, Pentose-phosphate, TCA cycle, ETC II/III/IV/V<br>Sulfate reduction ($SO_4^{2-} \rightarrow H_2S$)<br>Acetate, lactate and ethanol fermentation<br>Multi-haem *c*-type cytochromes (MHCs)<br>Peptidases (63, peptide and oligopeptide transporters, polar and branched AA transporters)<br>***C-fixation***<br>Calvin cycle, rTCA<br>***Other***<br>Flagellar assembly |
| | ERP019367_co2_29 | **Completeness:** 94%<br>**Contamination:** 3%<br>**Assembly size:** 6.5Mbp<br>**# contigs:** 155<br>**GC:** 39%<br>**rRNAs:** 5S<br>**tRNAs:** 15 | ***Source***<br>Groundwater<br>***Marker-gene***<br>Groundwater (2)<br>***Rarity***<br>Strictly moderate | ***Aerobicity***<br>Predicted anaerobic (99.9% confidence)<br>***Energy***<br>Glycolysis, Pentose-phosphate, TCA cycle, ETC I/II/III/IV/V<br>Sulfate reduction ($SO_4^{2-} \rightarrow H_2S$)<br>Acetate and propionate fermentation<br>Multi-haem *c*-type cytochromes (MHCs)<br>Peptidases (73, peptide transporter, polar and branched AA transporters)<br>***C-fixation***<br>Calvin cycle, rTCA<br>***Other***<br>Flagellar assembly |

| Phyla (habitats) | Genome | Stats | Habitat/s | Metabolism |
| --- | --- | --- | --- | --- |
| binchicken_b2<br><br>Rock (3) | SRP338002_co1_41 | **Completeness:** 94%<br>**Contamination:** 0.5%<br>**Assembly size:** 3.9Mbp<br>**# contigs:**<br>**GC:** 48%<br>**rRNAs:** 5S<br>**tRNAs:** 19 | ***Source***<br>Hydrothermal vents<br>***Marker-gene***<br>Rock (3)<br>***Rarity***<br>Conditionally rare | ***Aerobicity***<br>Predicted anaerobic (99.9% confidence)<br>***Energy***<br>Glycolysis, Pentose-phosphate, TCA cycle, ETC I/II/III/IV/V<br>Sulfate reduction ($SO_4^{2-} \rightarrow H_2S$)<br>Acetate, lactate, propionate and butyrate fermentation<br>Multi-haem *c*-type cytochromes (MHCs)<br>Peptidases (53, peptide transporter)<br>***C-fixation***<br>rTCA<br>***Other***<br>CRISPR-Cas Type I |
| binchicken_b3<br><br>Marine sediment (12)<br>Salt marsh (11)<br>Sediment (3)<br>Seawater (2)<br>Subsurface (2)<br>Hot springs (1) | binchicken_co34_576 | **Completeness:** 99%<br>**Contamination:** 4%<br>**Assembly size:** 3.8Mbp<br>**# contigs:** 349<br>**GC:** 44%<br>**rRNAs:** 5S, 16S, 23S<br>**tRNAs:** 19 | ***Source***<br>Salt marsh<br>***Marker-gene***<br>Salt marsh (11)<br>Marine sediment (5)<br>Aquatic subsurface (1)<br>***Rarity***<br>Strictly rare | ***Aerobicity***<br>Predicted anaerobic (99.9% confidence)<br>***Energy***<br>Glycolysis, Pentose-phosphate, ETC V<br>Homoacetogenesis (Wood-Ljungdahl)<br>Acetate, ethanol and butyrate fermentation<br>Peptidases (49, peptide transporter, polar and branched AA transporters)<br>***C-fixation***<br>Calvin cycle, rTCA, Homoacetogenesis (Wood-Ljungdahl)<br>***Other***<br>Amorphous cellulose cazyme |
| binchicken_b4<br><br>Groundwater (40)<br>Wastewater (17)<br>Freshwater sediment (12)<br>Wetland (5)<br>Activated sludge (5)<br>Sediment (5)<br>Freshwater (5)<br>Soil (4)<br>Bioreactor (4) | SRP089250_co1_180 | **Completeness:** 90%<br>**Contamination:** 3%<br>**Assembly size:** 3.6Mbp<br>**# contigs:** 367<br>**GC:** 65%<br>**rRNAs:** 5S<br>**tRNAs:** 17 | ***Source***<br>Activated sludge<br>***Marker-gene***<br>Activated sludge (2)<br>Freshwater (1)<br>Groundwater (1)<br>***Rarity***<br>Conditionally rare | ***Aerobicity***<br>Predicted anaerobic (99.9% confidence)<br>***Energy***<br>Glycolysis, Pentose-phosphate, ETC V<br>Sulfate reduction ($SO_4^{2-} \rightarrow SO_3^{2-}$)<br>Acetate, ethanol and lactate fermentation<br>Peptidases (47, peptide transporter, AA transporter)<br>***C-fixation***<br>Calvin cycle<br>***Other***<br>Flagellar assembly, Assimilatory sulfate reduction |

| **Phyla (habitats)** | **Genome** | **Stats** | **Habitat/s** | **Metabolism** |
|---|---|---|---|---|
| Subsurface (3)<br>Marine sediment (2)<br>Estuary (2)<br>Aquifer (1)<br>Hot springs (1)<br>Marine (1)<br>Biofilm (1)<br>Landfill (1)<br>Manure (1) | binchicken_co154_39 | **Completeness:** 90%<br>**Contamination:** 3.7%<br>**Assembly size:** 4.0Mbp<br>**# contigs:** 485<br>**GC:** 66%<br>**rRNAs:** 5S<br>**tRNAs:** 18 | ***Source***<br>Wastewater<br>***Marker-gene***<br>Wastewater (15)<br>Freshwater sediment (1)<br>***Rarity***<br>Strictly rare | ***Aerobicity***<br>Predicted anaerobic (99.9% confidence)<br>***Energy***<br>Glycolysis, Pentose-phosphate, ETC V<br>Sulfate reduction ($SO_4^{2-} \rightarrow SO_3^{2-}$)<br>Acetate, ethanol and lactate fermentation<br>Hydrogenase (type NiFe group 3d)<br>Peptidases (49, peptide transporter, branched AA transporter)<br>***C-fixation***<br>Calvin cycle<br>***Other***<br>Flagellar assembly, Assimilatory sulfate reduction |
| binchicken_b5<br><br>Sediment (18)<br>Activated sludge (10)<br>Wastewater (7)<br>Sludge (4)<br>Bioreactor (3)<br>Biofilm (3)<br>Biofilter (2)<br>Groundwater (1) | ERP124720_co1_46 | **Completeness:** 83%<br>**Contamination:** 2%<br>**Assembly size:** 3.4Mbp<br>**# contigs:** 672<br>**GC:** 61%<br>**rRNAs:** 5S<br>**tRNAs:** 19 | ***Source***<br>Groundwater (bioreactor)<br>***Marker-gene***<br>Sediment (14)<br>***Rarity***<br>Conditionally rare | ***Aerobicity***<br>Predicted aerobic (60% confidence)<br>***Energy***<br>Glycolysis, TCA cycle, Glyoxylate cycle, ETC V<br>Nitrate reduction ($NO_3^- \rightarrow NH_4^+$)<br>Acetate and lactate fermentation<br>Multi-haem *c*-type cytochromes (MHCs)<br>Peptidases (42, peptide transporter)<br>***C-fixation***<br>rTCA<br>***Other***<br>Amorphous cellulose cazyme, Flagellar assembly, glutamate → siroheme →heme (iron and heme transporters) |
| | binchicken_co350_29 | **Completeness:** 69%<br>**Contamination:** 1%<br>**Assembly size:** 3.0Mbp<br>**# contigs:** 841<br>**GC:** 65%<br>**rRNAs:** -<br>**tRNAs:** 16 | ***Source***<br>Activated sludge<br>***Marker-gene***<br>Activated sludge (2)<br>***Rarity***<br>Strictly rare | ***Aerobicity***<br>Predicted anaerobic (53% confidence)<br>***Energy***<br>TCA cycle, ETC II/V<br>Nitrate reduction ($NO_3^- \rightarrow NH_4^+$)<br>Acetate and lactate fermentation<br>Multi-haem *c*-type cytochromes (MHCs)<br>Peptidases (39, peptide transporter, branched AA transporter)<br>***C-fixation***<br>rTCA<br>***Other***<br>Flagellar assembly |

| **Phyla (habitats)** | **Genome** | **Stats** | **Habitat/s** | **Metabolism** |
|---|---|---|---|---|
| binchicken_b6<br><br>Marine (45)<br>Seawater (12)<br>Cold seep (7)<br>Salt lake (6)<br>Marine sediment (6)<br>Sediment (4)<br>Hydrothermal vent (2)<br>Aquatic (1)<br>Estuary (1) | SRP077602_co1_83 | **Completeness:** 95%<br>**Contamination:** 6%<br>**Assembly size:** 5.7Mbp<br>**# contigs:** 202<br>**GC:** 54%<br>**rRNAs:** 5S<br>**tRNAs:** 18 | ***Source***<br>Marine<br>***Marker-gene***<br>Marine (14)<br>Seawater (6)<br>***Rarity***<br>Conditionally rare | ***Aerobicity***<br>Predicted anaerobic (99.9% confidence)<br>***Energy***<br>Glycolysis, Pentose-phosphate, TCA cycle, ETC II/V<br>Sulfate reduction ($SO_4^{2-} \rightarrow SO_3^{2-}$)<br>Acetate, ethanol and butyrate fermentation<br>Hydrogenases (2 type NiFe group 3c)<br>Peptidases (61, peptide and oligopeptide transporters)<br>***C-fixation***<br>Calvin cycle, rTCA<br>***Other***<br>Flagellar assembly, Assimilatory sulfate reduction |
| | SRP308112_co1_304 | **Completeness:** 91%<br>**Contamination:** 4%<br>**Assembly size:** 4.4Mbp<br>**# contigs:** 82<br>**GC:** 52%<br>**rRNAs:** 5S, 16S<br>**tRNAs:** 17 | ***Source***<br>Marine<br>***Marker-gene***<br>Marine (28)<br>Seawater (5)<br>***Rarity***<br>Conditionally rare | ***Aerobicity***<br>Predicted anaerobic (99.9% confidence)<br>***Energy***<br>Glycolysis, Pentose-phosphate, ETC V<br>Nitrate reduction (napA + narH)<br>Acetate, ethanol and butyrate fermentation<br>Peptidases (52, peptide and oligopeptide transporters)<br>***C-fixation***<br>Calvin cycle, rTCA<br>***Other***<br>Flagellar assembly, Assimilatory sulfate reduction |
